# Structure and function of otoferlin, a synaptic protein of sensory hair cells essential for hearing

**DOI:** 10.1101/2025.05.10.653085

**Authors:** Han Chen, Constantin Cretu, Abigail Trebilcock, Natalia Evdokimova, Norbert Babai, Laura Feldmann, Florian Leidner, Fritz Benseler, Sophia Mutschall, Klara Esch, Csaba Zoltan Kibedi Szabo, Vladimir Pena, Constantin Pape, Helmut Grubmüller, Nicola Strenzke, Nils Brose, Carolin Wichmann, Julia Preobraschenski, Tobias Moser

## Abstract

Our sense of hearing relies upon speedy synaptic transmission of sound information from cochlear inner hair cells (IHCs) to spiral ganglion neurons (SGNs). To accomplish this, IHCs employ a sophisticated presynaptic machinery including the multi-C_2_-domain protein otoferlin which is affected by human deafness mutations. Otoferlin is essential for IHC-exocytosis but how it binds Ca^2+^ and the target membrane to serve synaptic vesicle (SV) tethering, docking and fusion remained unclear. Here, we obtained cryo-electron-microscopy structures of Ca^2+^-bound otoferlin and employed molecular dynamics simulations of membrane binding. We show that membrane binding involves C_2_B-C_2_G-domains and repositions C_2_F- and C_2_G-domains. Progressive disruption of Ca^2+^-binding by the C_2_D-domain in mice increasingly altered synaptic sound encoding and eliminated the Ca^2+^-cooperativity of SV-exocytosis, indicating that this Ca^2+^-cooperativity reflects binding of several Ca^2+^-ions to otoferlin. Together, our findings elucidate molecular mechanisms underlying otoferlin-mediated SV-docking and support a role of otoferlin as Ca^2+^-sensor of SV-fusion in IHCs.

## Introduction

Inner hair cells (IHCs) and spiral ganglion neurons (SGNs) form highly specialized ribbon synapses to achieve indefatigable sound encoding at spike rates of hundreds per second and with sub-millisecond temporal precision (*1–3*). These high functional demands of reliably releasing SVs at even greater rates in synchronicity with the stimulus are thought to have shaped an unconventional molecular synaptic machinery that deviates from synapses of the central nervous system (CNS, (e.g. *4–13*)). However, the composition and function of this machinery has remained elusive. For example, the identity of the Ca^2+^-sensor of SV-fusion and the role of soluble *N*-ethylmaleimide-sensitive factor attachment protein receptors (SNARE) proteins and SNARE-regulators at the IHC-SGN synapse are currently unknown (*11–14*).

Human deafness mutations leading to auditory synaptopathy (*15*) affect at least two key players of the IHC synaptic machinery, Ca_V_1.3 Ca^2+^-channel complexes and otoferlin (*16–18*). Ca_V_1.3 Ca^2+^-channels couple the IHC receptor potential to SV-exocytosis. They activate at low voltages, inactivate slowly and are positioned in nanometer proximity to SV release sites for tight control of glutamate release (*19–22*). Otoferlin is a Ca^2+^-sensitive multi-C_2_ domain protein that is thought to reside on both SVs and the active zone (AZ) membrane of IHCs (*6*, *23*, *24*). Otoferlin comprises at least seven C_2_ domains (C_2_A-C_2_G) of which four (C_2_C, C_2_D, C_2_F and C_2_G) are thought to bind Ca^2+^, involving aspartate residues in their top loops and likely also phospholipids in target membrane (ref. (*6*, *25*, *26*)). In analogy to synaptotagmins 1 or 2 at CNS synapses (*27*, *28*), otoferlin is thought to act as the Ca^2+^-sensor of SV-fusion in IHC (*6*, *25*, *26*, *29*). Physiological data indicate that the cooperative binding of 4-5 Ca^2+^ ions to the Ca^2+^-sensor (“Ca^2+^-cooperativity”) is required for SV-fusion to occur in IHCs (*22*, *30*).

Analysis of mouse mutants for *Otof* and interacting proteins has indicated a multifaceted role of otoferlin in the SV-cycle of IHCs. Beyond its putative role as Ca^2+^-sensor of SV-fusion, otoferlin is required for SV replenishment to the readily releasable pool (RRP, (*12*, *24*, *26*, *31*, *32*)) likely acting as an SV-tethering and -docking factor (*12*), and for coupling exo- and endocytosis (*33–35*). However, even for the first postulated role as Ca^2+^-sensor function of otoferlin in SV-fusion, unequivocal demonstration has remained challenging. This is due to i) difficulties to disentangle otoferlin’s action in Ca^2+^-triggered SV-fusion vs. Ca^2+^-dependent SV-replenishment (*26*, *31*), ii) the fact that IHC exocytosis is largely abolished in mice with mutations that target Ca^2+^-binding to C_2_F (C_2_E in previous nomenclature, (*29*)) and C_2_G (i.e. C_2_F, (*36*)), which precludes rigorous testing of the Ca^2+^-sensor of SV-fusion hypothesis and iii) challenges in purifying sufficient amounts of full-length protein for *in vitro* studies of otoferlin structure and function (*29*, *37*). So far, to our knowledge, only a crystal structure of the C_2_A domain (*38*), the AlphaFold structure prediction and optical nanoscopy images of otoferlin (*39*) are available in this regard.

The paucity of structural data on otoferlin in its lipid-free and membrane-bound state has limited our understanding of the molecular mechanisms at the basis of otoferlin function. In particular, we do not know how individual C_2_ domains bind Ca^2+^ and phospholipids and how they participate in SV-tethering, -docking and -fusion. Progress in understanding the structure and function of otoferlin is urgently needed, given that its defects cause deafness in humans with hundreds of missense mutations that have remained hard if not impossible to interpret. Moreover, only a few years after preclinical proof of concept (*40*, *41*), recent clinical gene therapy trials to remedy otoferlin-related deafness have yielded first promising results (*42–44*) and detailed information on the otoferlin structure and function is needed to optimize these translational approaches.

We took a multidisciplinary approach to elucidate the mechanisms by which otoferlin operates. We combined *in vitro* work on the structure and function of purified otoferlin with analyses of *Otof* mouse mutants. Using cryo-electron-microscopy (cryo-EM) and molecular dynamics simulation, we reveal how otoferlin engages lipid membranes with prominent roles of the N-terminal C_2_B and the Ca^2+^-bound C-terminal C_2_F and C_2_G domains that are essential for SV-exocytosis (*29*, *35*, *36*). Membrane binding by C_2_F and C_2_G is accompanied by their repositioning likely mediating SV-docking at the AZ. Ca^2+^-binding by C_2_G is required for otoferlin to efficiently bind the target membrane *en route* to vesicle fusion. Multiscale physiology of sound encoding in the *Otof* mouse mutants with structure-guided progressive alteration of Ca^2+^-binding to the C_2_D domain revealed increasingly reduced SV-release probability but spared SV-replenishment at the IHC AZ. Probing the apparent Ca^2+^-dependence of IHC exocytosis revealed a loss of the physiological Ca^2+^-cooperativity indicating that otoferlin serves as Ca^2+^-sensor of SV-fusion in IHCs.

## Results

### Cryo-EM structures of mouse otoferlin in the membrane-bound and lipid-free states

We engineered a soluble variant of mouse otoferlin (residues 216-1931) for structural studies (Fig. 1A and Fig. S1A). The construct comprised all the cytosolic domains of otoferlin, except for the non-essential (*45*) N-terminal C_2_A, which has not been linked to known missense mutations in patients (*46*), and lacked the C-terminal transmembrane helix (Fig. 1A). The purified otoferlin sample was well-folded (Fig. S1B-C), retained Ca^2+^-binding activity (Fig. S1D) and stably interacted in a Ca^2+^-dependent manner with model lipid membranes (liposomes) containing the anionic phospholipids phosphatidylserine (PS) and PI(4,5)P_2_ (phosphatidyl inositol-4,5-bisphosphate) (Fig. S1E and S1G). Otoferlin showed stronger preference for Ca^2+^ over Mg^2+^ in lipid binding than myoferlin (*47*) (Fig. S1G). Size-exclusion chromatography and mass photometry of soluble otoferlin (216–1931) predominantly revealed monomeric species (Fig. S1B-C and S1F). To obtain insights into the organization of otoferlin and its mode of lipid membranes interaction, we obtained Ca^2+^-bound cryo-EM structures of otoferlin (216–1931) in the lipid-free (Fig. 1B-C and Fig. S3A-C) and membrane-bound states (Fig. 1D-E and Fig. S2A-E), resolved at a global resolution of ∼2.9 Å and ∼2.3 Å, respectively (Table S1). To obtain the membrane-bound structure we employed membrane scaffold protein 2 N2 (MSP2N2)-based nanodiscs as lipid bilayer surrogates (*48*) (Fig. S2A-C). Pre-assembled MSP2N2 lipid nanodiscs containing anionic phospholipids (30 mol% PS and 10 mol% PI(4,5)P_2_) formed stable complexes with otoferlin (216–1931) in the presence of Ca^2+^. This stabilized the structure, allowing for the accurate modeling of the C_2_B-C_2_G domains, the Fer domain, the linker regions, multiple Ca^2+^ ions and phospholipid-binding sites (Fig. 1D-E, Fig. S2C-H and Fig. S3D). Conversely, in the absence of lipid membranes (Fig. 1B-C and Fig. S3A-C), otoferlin exhibited increased dynamics of the C-terminal C_2_G and required additional computational sorting to identify discrete conformational states (Fig. S3A-C). The lipid-bound otoferlin adopts a compact tertiary structure, reminiscent of a closed ring (Fig. 1D-E, Fig. S2C, Movie S1) which spans ∼128 Å by ∼98 Å and bears similarity to models of lipid-bound myoferlin (*47*). The individual C_2_ domains are distributed sequentially, starting from the N-terminal C_2_B to the C-terminal C_2_G domains, around the cavity of the ferlin ring. The well-resolved C_2_C and C_2_D domains form the top arch of the ring which is rigidified through the C_2_CD-FerA module (Fig. 1D and Fig. S2C), generally organized as in other ferlin structures (*47*, *49*). C_2_B and C_2_E occupy diametrically opposed locations on the ring and establish interfaces with C_2_C and C_2_D, respectively, and with the C-terminal C_2_F and C_2_G (Fig. 1E). The tertiary structure of otoferlin in the lipid-bound state appears to be further reinforced by several accessory structural motifs and linker regions (Fig. 1E).

**Figure 1.**
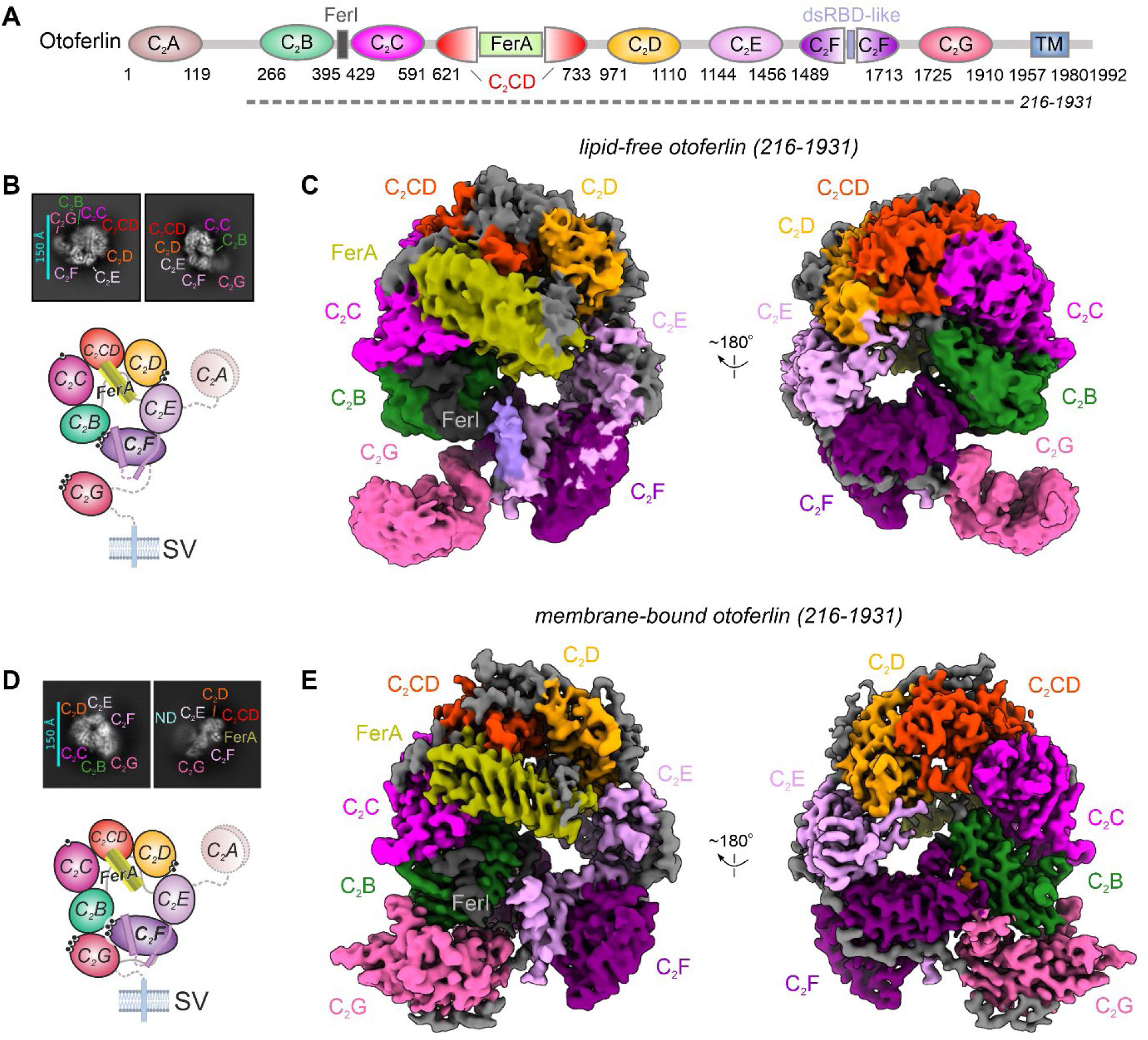
Cryo-EM structures of mouse otoferlin in the lipid-free and membrane-bound states. **(A)** Schematic domain composition of mouse otoferlin based on the structure. The structural motifs of otoferlin are depicted as coloured circles or boxes (domain colour scheme used throughout the manuscript), whereas the FerI motif and linker regions are coloured in grey. The borders of the otoferlin construct (residues 216-1931) used for structural studies are indicated by the dashed line. **(B)** Selected 2D classes averages of otoferlin (216–1931), in the lipid-free state. The assigned C_2_ domains are indicated and also shown in the cartoon representation in the bottom panel, with a hypothetical position of the missing C_2_A domain. **(C)** Overall cryo-EM map of otoferlin (residues 216-1931) in the lipid-free state. The otoferlin cryo-EM map has been coloured after the modelled domains and is shown in two different orientations. **(D)** 2D class averages and structure schematic of mouse otoferlin in the membrane-bound state. The MSP2N2 nanodisc (ND, cyan) density is indicated together with the visible structural motifs. **(E)** The cryo-EM map of otoferlin (residues 216-1931), resolved in the membrane(nanodisc)-bound state. The map has been rendered as in **(C).**

The C_2_A-C_2_B linker (residues 241-265), connecting the N-terminal C_2_A (absent from the structure) to C_2_B, adopts an extended conformation, appearing to wrap around the FerA four-helix bundle (residues 736-853) and C_2_D before reaching, on the other side of the ring, the membrane-bound C_2_B. In addition, the conserved FerI motif (residues 396-428), which connects C_2_B and C_2_C, interacts with both C_2_F and C_2_B while also stabilizing the C-terminal C_2_G (Fig. 1E). C_2_G samples the conformational space between C_2_B and C_2_F, like in myoferlin and dysferlin structures (*47*), and is poorly resolved in the lipid-free state of otoferlin (Fig. 1B-C). Overall, this structural feature might indicate that the C_2_G interfaces with FerI and the neighboring C_2_B and C_2_F are established and stabilized through membrane binding.

### Membrane binding interfaces of otoferlin

In our cryo-EM structure, otoferlin engages the lipid (nanodisc) bilayer asymmetrically through a multi-domain interface formed by the N-terminal C_2_B and the C-terminal C_2_F-C_2_G (Fig. 2A, 2C and 2E, Movie S2). The three domains are positioned close to each other on the nanodisc and their top loops (L1 and L3) project on one side to deeply insert into the lipid membrane. As a result of these interactions, the ferlin ring appears tilted by ∼40° with respect to the membrane plane. The N-terminal C_2_B lacks the characteristic L1-L3 aspartate residues required for Ca^2+^-coordination as also found in other ferlins (*47*). Consequently, C_2_B binds the membrane via its concave side through multiple residues positioned at the tip of the long L3 loop independent of Ca^2+^-coordination (Fig. 2B). Interestingly, the hydrophobic residues (L341 and L342) of C_2_B appear to penetrate and thereby to modulate the nanodisc bilayer, whereas the interacting basic residues (R343 and K339) establish more superficial contacts at the surface, possibly with phospholipid headgroups (Fig. 2B). In contrast, both C_2_F and C_2_G (Fig. 2D and Fig. 2F) are oriented towards the nanodisc head-on allowing for the aspartate residues of L1 and L3 to engage the lipid bilayer directly in a Ca^2+^-sensitive manner, consistent with the domains playing a key role in late-stage SV exocytosis (*29*, *36*).

**Figure 2.**
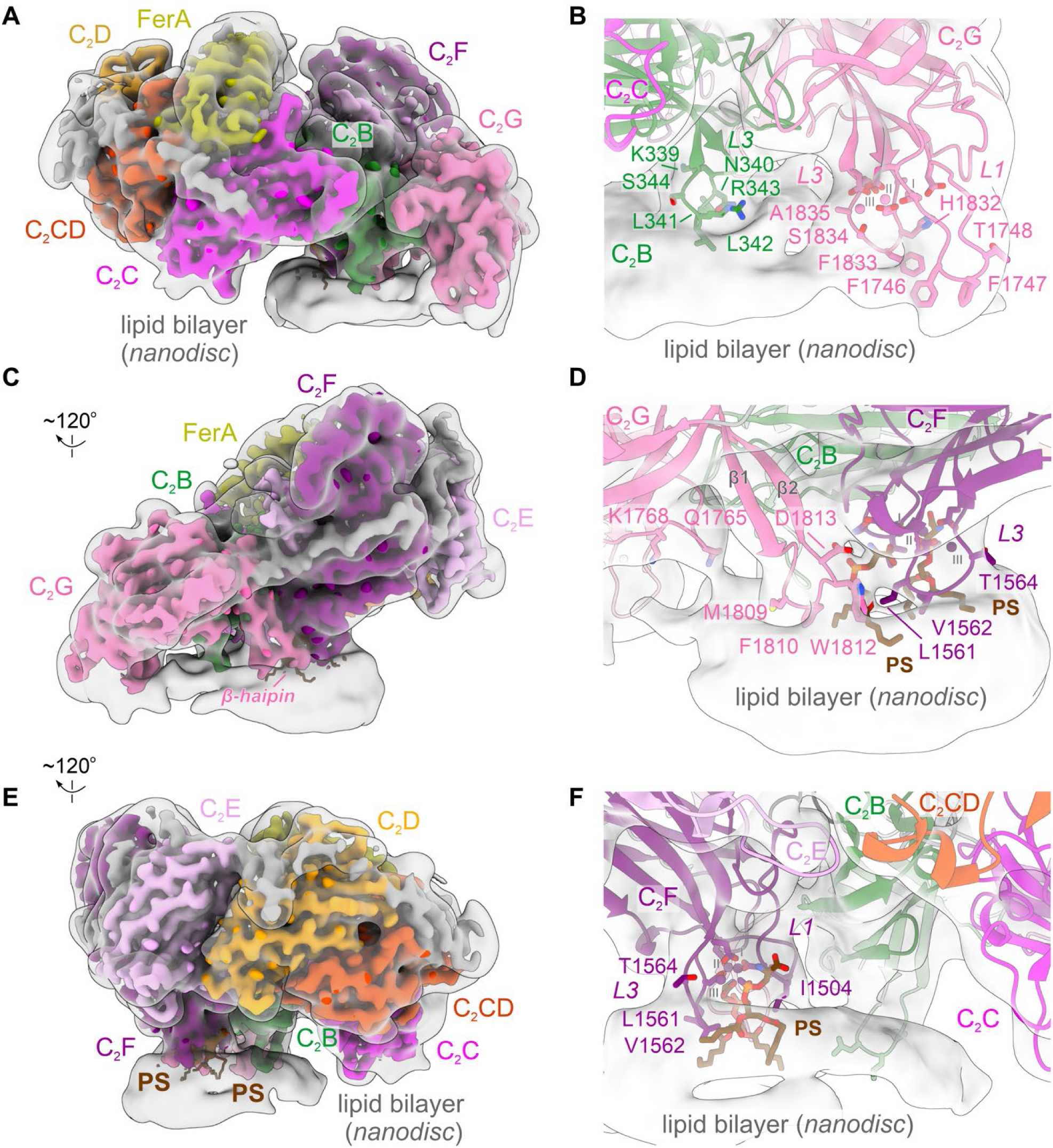
The multi-domain membrane-binding interface of otoferlin. **(A)** The high-resolution cryo-EM map of mouse otoferlin (216–1931) bound to a lipid nanodisc and viewed from its N-terminal side. The map has been colored as in Fig. 1E and, to enable the visualization of the ordered nanodisc regions, a transparent, low-pass filtered (to 8 Å) map of the otoferlin-nanodisc complex has been superimposed onto the high-resolution map. **(B)** The lipid nanodisc contact interfaces of the C_2_B and C_2_G domains, oriented as in A. The otoferlin model is depicted as cartoon. The nanodisc density is shown as a low-pass filtered (to 8 Å) cryo-EM map. The interface and Ca^2+^-coordinating residues are shown as sticks. The modelled Ca^2+^-binding sites are indicated with roman numerals (I-III). **(C)** The composite lipid binding interface formed by the C_2_F and C_2_G domains. The otoferlin maps are displayed as in **(A)**. **(D)** Close-up view of the membrane interacting β-hairpin of C_2_G and the L3 loop of C_2_F. The otoferlin residues projecting towards the surface of the lipid nanodisc are shown as sticks and the model is fitted inside the low-passed (to 8 Å) map of the otoferlin-nanodisc complex. The two recruited phosphatidylserine (PS) headgroups are depicted as sticks and colored brown. **(E)** C_2_C, C_2_D and C_2_E do not engage the lipid membrane in the otoferlin-nanodisc complex. The cryo-EM otoferlin-nanodisc maps are rendered and colored as in **(A)**. **(F)** C_2_F recruits two PS headgroups, originating from the nanodisc bilayer, to its three Ca^2+^-binding sites. The lipid interface residues are depicted as sticks and the PS molecules are colored brown. See also **(D)**.

Importantly, in both cases the bound Ca^2+^ ions form pockets for interaction with phospholipids and likely orient the L1 (C_2_G) and L3 (C_2_F and C_2_G) loop residues to facilitate their membrane insertion (Fig. 2D and Fig. 2F), resulting in a complex network of membrane contacts. Apart from the Ca^2+^-facilitated membrane contacts, C_2_F and C_2_G form a likely lipid-induced tertiary interface between the β-hairpin motif of C_2_G and the L4 loop of C_2_F. In addition, the β-hairpin of C_2_G itself binds the lipid membrane through several hydrophobic residues (W1812, F1810, M1809), while also framing the phospholipid (PS) binding pocket of C_2_F (Fig. 2D).

### Ca^2+^ and lipid binding sites of otoferlin

Biophysical studies of isolated C_2_ domains of otoferlin estimated that 2-3 Ca^2+^ ions are bound per domain and that the domains have different divalent-specificity and Ca^2+^-sensitivity for binding the acidic phospholipids PS and PI(4,5)P_2_ (*25*, *50*, *51*). We could model nine Ca^2+^ ions (C_2_C: 1, C_2_D: 2, C_2_F: 3 and C_2_G: 3) of which six were part of the lipid interface of otoferlin (**Fig. 3A-C and Fig. 3E**) and two PS molecules of the nanodisc contributing to the coordination of the three Ca^2+^ by the C_2_F domain (Fig. 3D). In contrast to previous reports (*25*, *50*), the cryo-EM structure showed that C_2_B and C_2_E are not involved in Ca^2+^-coordination. However, the structural data confirmed the predicted Ca^2+^-coordination by top loop aspartates D1558 (D1563 in human) and D1560 (D1565 in human) of C_2_F. These were replaced by alanine to disrupt Ca^2+^-binding in the *Otof^TDA^* mouse mutant, which abolished Ca^2+^-influx-triggered exocytosis in IHCs despite sizable (∼60%) otoferlin levels (*29*). Here, we performed electron tomography of IHC synapses in conventionally embedded organs of Corti of *Otof^TDA^*mice and did not find an obvious SV-docking deficit (Fig. S4). This might suggest that TDA-otoferlin retains sufficient membrane binding activity. However, we cannot exclude a deficit in tight SV-docking that could be essential for SVs to gain full fusion competence (*52*). C_2_F Ca^2+^-coordination further involves D1509, A1502 (backbone carbonyl) and D1503 residues of L1 and D1558 and W1559 (backbone carbonyl) residues of L3 (Fig. 3C). The two PS headgroups bound by C_2_F complete the coordination shells of the first and third Ca^2+^ sites, where they interact through the serine or glycerophosphate moieties, respectively. The side chains of the two bound PS are only partially embedded into the lipid nanodisc, suggesting that they were “pulled” from the nanodisc bilayer by C_2_F (Fig. 3D, F). Indeed, the PS at the first Ca^2+^ site appears to establish additional hydrophobic interactions with residues F1810 and W1812 of C_2_G’s β-hairpin that line the binding pocket and, likely, stabilize the bound PS. This dual PS (Fig. 3F) recognition mechanism employed by otoferlin’s C_2_F appears unique and likely evolved to allow for a high degree of membrane insertion and local remodelling (Fig. 3D, F).

**Figure 3.**
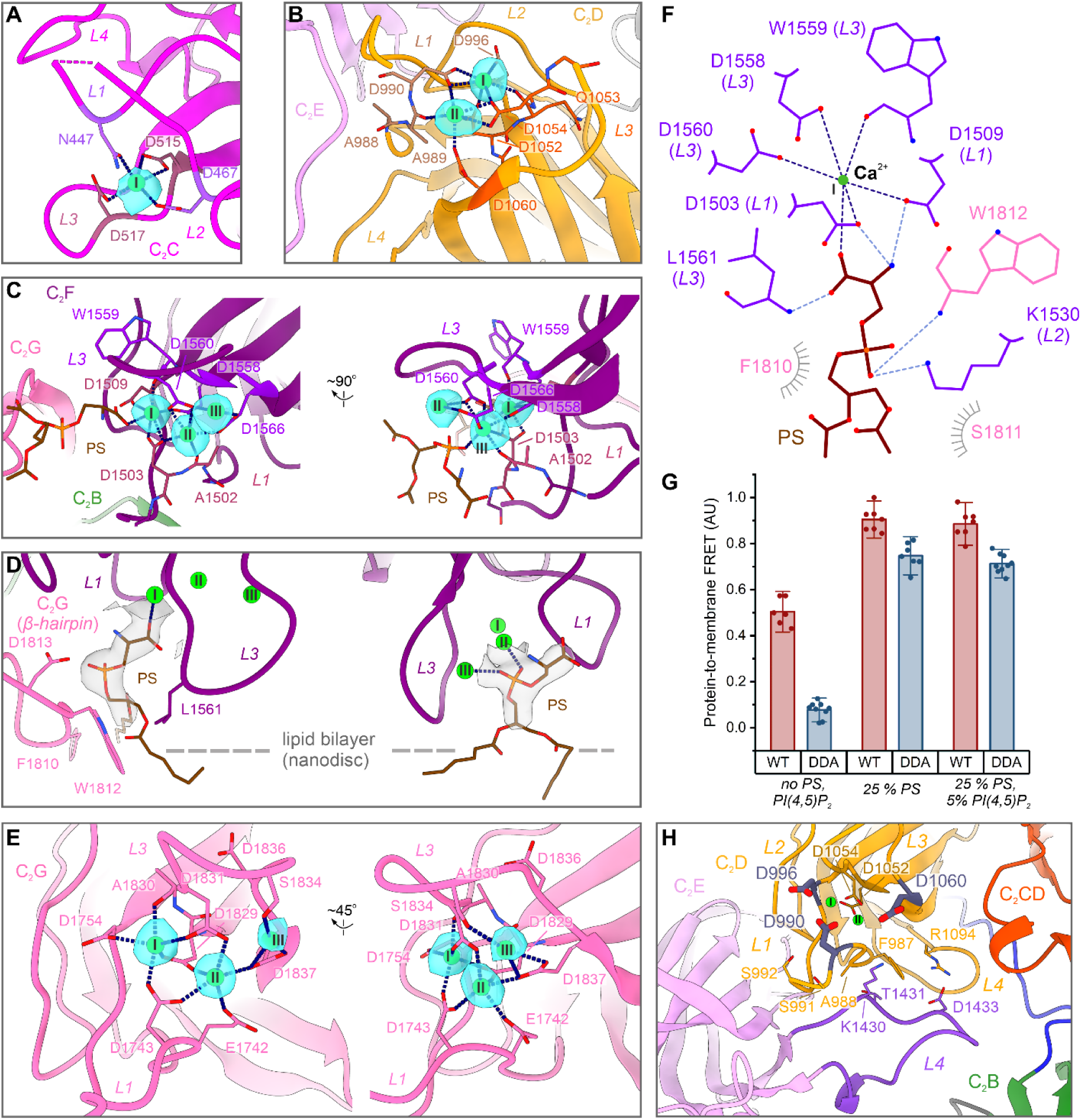
Ca^2+^ and lipid-binding sites modelled in otoferlin’s membrane-bound structure. **(A)** Ca^2+^-binding by the C_2_C domain. Residues forming the Ca^2+^-binding pocket are depicted as sticks. The normalized cryo-EM difference map (cyan, 2σ) is contoured around the modelled Ca^2+^ ion. **(B)** The two Ca^2+^ sites of the C_2_D domain. The cryo-EM difference map of otoferlin-nanodisc complex (cyan, 5σ) is contoured around the two modelled Ca^2+^ ions. The Ca^2+^-binding residues of C_2_D are shown as sticks. **(C)** The modelled Ca^2+^ ions of otoferlin’s C_2_F domain. The three Ca^2+^ sites are shown in two different orientations and the cryo-EM difference map (cyan, 5σ) is displayed around the modelled ions. The two bound PS molecules are shown as sticks and coloured brown. **(D)** C_2_F binds two PS molecules, originating from the lipid nanodisc bilayer, through three coordinated Ca^2+^ ions. The cryo-EM density map of the otoferlin-nanodisc complex (grey surface) is contoured around the modelled ligands. The nanodisc surface is indicated schematically through a dashed line. **(E)** C_2_G coordinates three Ca^2+^ ions in the membrane-bound state of otoferlin. The structural model and the normalized cryo-EM difference map (cyan, 5σ) are displayed in two orientations. **(F)** Schematic depicting the unique PS headgroup recognition mechanism employed by the C_2_F domain of otoferlin. Note that the seryl moiety interacts with the coordinated Ca^2+^ ion (at the 1^st^ Ca^2+^-binding site, I), two aspartate residues (D1503, D1509) and a leucine (L1561), lining the pocket. The Ca^2+^-coordination bonds are shown as dashed dark blue lines, whereas the polar contacts to the glycerophosphate moiety are coloured in light blue. The hydrophobic interactions are indicated as grey half-circles. **(G)** The liposome-binding activity of otoferlin (216–1931)-D1829A/D1831A (DDA) mutant of the C_2_G domain, harbouring two neutralizing substitutions (“Double Aspartate (D) to Alanine (A)”). For direct comparison, the wild-type (WT) protein was analysed in parallel. The FRET-based measurements were carried out in the presence of 250 µM CaCl_2_ and at 0.25 µM and 50 µM protein and liposome concentrations, respectively. The measurements were repeated n=6 (WT, in the presence of liposomes lacking PS and PI(4,5)P_2_, denoted as no PS, PI(4,5)P_2_), n=9 (DDA, in the presence of liposomes comprising 25 mol% PS and 5 mol% PI(4,5)P_2_ or lacking PS and PI(4,5)P_2_), or n=7 (remaining measurements). The data were normalized to the maximum Dansyl-PE emission after subtraction of the background from each individual point (using protein-free measurements) and error bars indicate the standard deviation (SD). See also Fig. S1G. **(H)** The tertiary interface between C_2_D and the extended L4 loop of C_2_E, which might be sensitive to Ca^2+^. C_2_D residues substituted in the mouse models (see also Fig. 6) are indicated in dark blue. The two Ca^2+^ sites of C_2_D are shown as spheres and coloured green.

Like C_2_F, C_2_G coordinates three Ca^2+^ ions through residues of the L1 (E1742, D1743, D1754) and L3 (D1829, A1830 (backbone carbonyl), D1831, S1834 and D1837) loops (Fig. 3E). D1837 (along with D1836) was substituted by alanine in *Otof^DDA^* mice (*36*) that showed reduced (∼60%) basolateral otoferlin levels in IHC, abolished IHC exocytosis, and deafness. Moreover, D1842 (human equivalent of mouse D1837) is a target of a human *OTOF* variant (*53*) (Fig. 8A, B). The top loops of C_2_G are rich in aromatic residues (F1746 (L1), F1747 (L1), F1833 (L3), Y1775 (L2)) and histidines (H1776 (L2), H1832 (L2)) which all appear to either insert into the membrane or position close to it. This might suggest that membrane binding involves a Ca^2+^-independent component. As our FRET-based assay (Fig. 3G and Fig. S1G) showed a decrease in liposome binding for the D1829A/D1831A mutant that targets two major Ca^2+^-binding sites of C_2_G, Ca^2+^-binding by C_2_G might still be needed for efficient lipid binding which is consistent with the strong exocytosis deficit in *Otof^DDA^*IHCs (*36*). Potentially, Ca^2+^-binding to C_2_G could rigidify and, consequently, help orient the long top loops of the domain towards the membrane plane – alternative to a direct phospholipid interaction mechanism.

Different from C_2_F and C_2_G, the Ca^2+^-binding top loops of C_2_C and C_2_D in the membrane-bound otoferlin structure point away from the nanodisc-membrane (Fig. 2E and Fig. 3A-B, but see results of molecular dynamics simulation below). Our structure shows that Ca^2+^-binding to C_2_C involves D467 and N447 residues of L1 and D515 and D517 residues of L3, targeted by *Otof^D515-517A^* mice (*26*), for which changes in the Ca^2+^-dependencies of SV-fusion and -replenishment were reported. Finally, C_2_D domain coordinates two Ca^2+^ ions mainly via the top loop aspartates: D990, D996 (L1), D1052 and Q1053 (backbone carbonyl) (L3) form the first Ca^2+^-binding site, while A989 (backbone carbonyl), D990 (LL1) as well as D1054 and D1060 (L3) coordinate the second Ca^2+^ ion (Fig. 3B, H). Interestingly, L1 of C_2_D is closely positioned to the extended L4 of the adjacent C_2_E (Fig. 3H). This represents the only tertiary interface of otoferlin which might be sensitive to Ca^2+^-binding, raising the possibility that C_2_D Ca^2+^-binding could serve a structural role and modulate the conformational states of the ferlin ring during the SV fusion cycle (see also Fig. 10).

### Conformational changes of otoferlin upon membrane binding

Whereas the top arch of the ferlin ring (C_2_B-C_2_D) is similar in the lipid-free and membrane-bound structures, the C-terminal C_2_F and C_2_G of membrane-bound otoferlin are rearranged generating a more compact, ‘closed’ conformational state (see also Movie S3). Comparison between the lipid-free ‘open’ state and the nanodisc-bound ‘closed’ state (see also Movie S4), indicates C_2_F moved locally by ∼9 Å in the direction of C_2_B and C_2_E. At the same time, the C_2_G domain, which samples multiple conformations in the lipid-free state, underwent a large-scale displacement by ∼21-34 Å towards the nanodisc plane (Fig. S3C and S3E). In its new pose, C_2_G established new contact interfaces, absent in the lipid-free state, with the convex surface of C_2_B (Fig. 4B), the FerI motif (Fig. 4C) and C_2_F via the β-hairpin (Fig. 4D), tailing a total ∼901 Å^2^. The combined domain movements appear to facilitate the in-plane positioning of the membrane-facing top loops of C_2_B (L3), C_2_F (L3) and C_2_G (L1 and L3) and allow their cooperative interaction with the nanodisc through a multi-domain binding interface. The formation of the new tertiary contacts, involving C_2_G, appears to also rigidify the overall structure and fix the, otherwise dynamic, C_2_B-C_2_F and C_2_E-C_2_F interfaces (Fig. S3E), further promoting stable membrane binding.

**Figure 4.**
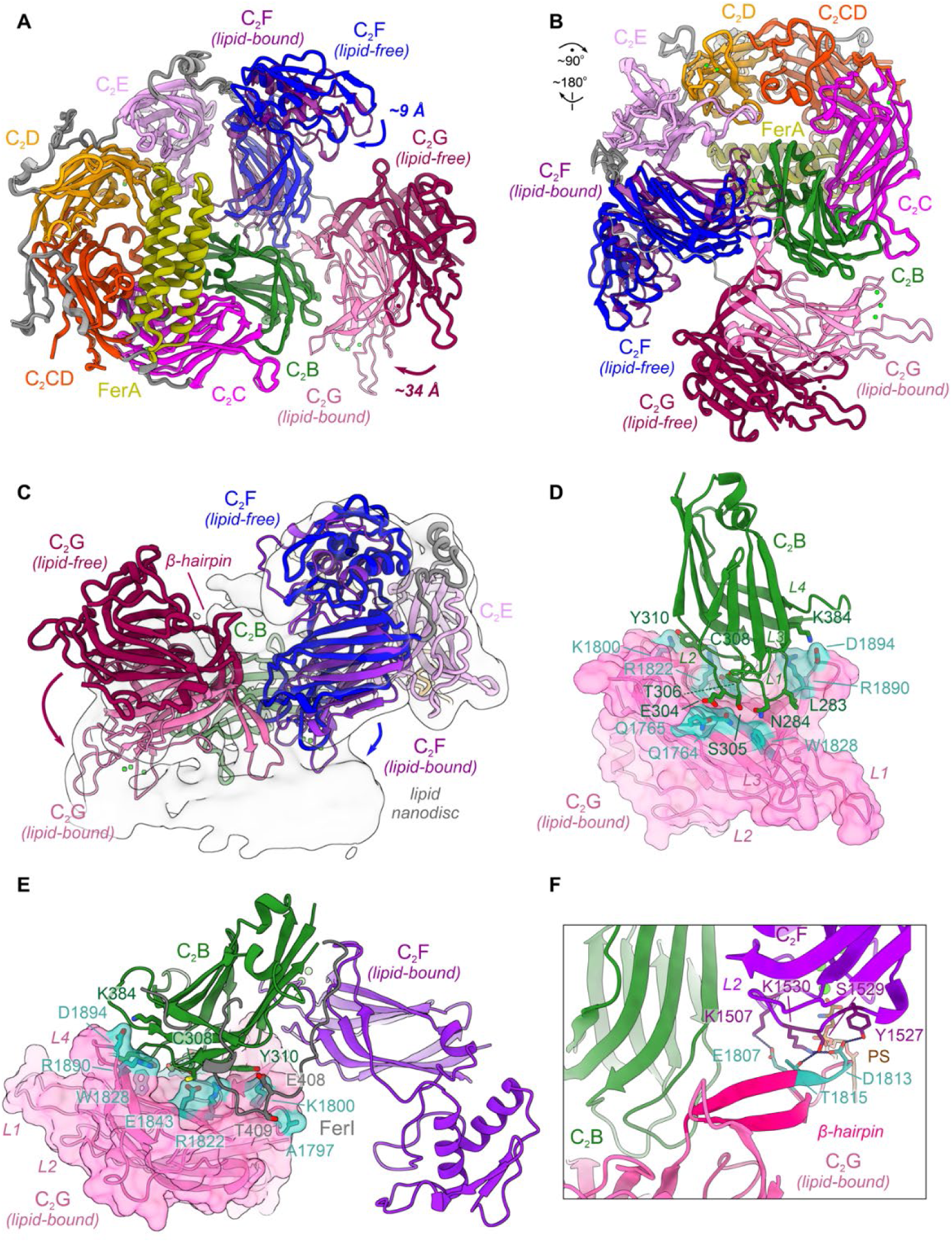
Conformational states of otoferlin. **(A)** Structural superposition between otoferlin (216-1931) in the lipid-free (‘open’) and nanodisc-bound (‘closed’) states. The otoferlin model in the closed state is depicted as a transparent cartoon representation, visualized from the N-terminal side. **(B)** The C_2_F (blue) and C_2_G (dark red) domains undergo a large-scale repositioning upon binding to a nanodisc bilayer. The superimposed lipid-free and lipid-bound models of otoferlin are shown from the membrane-binding side. **(C)** The repositioned C_2_F and C_2_G domains in the closed state are engaged in extended contacts with the lipid nanodisc. The otoferlin models in the closed and open states are shown together with a low-pass filtered map (to 8 Å) of the otoferlin-nanodisc complex. The C_2_F and C_2_G domains in the lipid-free state are coloured blue and dark red, respectively, and their movement upon nanodisc binding is indicated by arrows. **(D)** The new tertiary interface between the C_2_B and C_2_G domains established in the closed (membrane-bound) state of otoferlin. The C_2_B and C_2_G residues engaged in both polar and hydrophobic interactions (within 3.5 Å) are depicted as sticks. C_2_G is shown as a transparent solvent excluded surface. **(E)** The contact interfaces between C_2_B, C_2_G and the FerI motif in the closed state of otoferlin. Interacting residues within a 3.5 Å distance are depicted as sticks. The FerI motif is coloured grey. C_2_G is shown as a transparent solvent excluded surface. **(F)** Polar contacts between the β-hairpin motif of otoferlin and the top loop L2 of C_2_F observed in the lipid-bound state of otoferlin. The observed polar contacts are depicted as dashed blue lines. The neighboring C_2_B is coloured green.

Compared to myoferlin structures (*47*), the observed conformational transition of otoferlin is less complex in nature and, except C_2_G, all interacting domains are posed to engage the membrane, without requiring significant rearrangement from the lipid-free state. It is, therefore, possible that the pre-positioning of C_2_B, likely through the specific C_2_A-C_2_B linker region (residues 241-265), and, to a large extent, of C_2_F has evolved to support the faster timescales of otoferlin-dependent SV-exocytosis in hair cells (*54–56*) (milliseconds, compared to slower dysferlin/myoferlin-dependent fusion events occurring in tens of seconds (*57*)). Likely, for a similar reason, the membrane-proximal C_2_G is highly dynamic in the lipid-free state, a feature which might increase the target membrane capture radius of otoferlin to promote both fast SV-docking and -undocking, which we postulate to support the high rates of otoferlin-dependent SV-replenishment. Further supporting this hypothesis, computational analysis of otoferlin’s conformational space in the lipid-free state indicates that ‘closed-like’ conformations are sampled in the absence of a membrane, together with additional, likely, less stable, intermediate states. In these states, the C_2_G domain occupies both peripheral positions (farther away from C_2_B), as well as appearing to transiently engage C_2_F and closely approach the N-terminal C_2_B (Fig. S3C and S3E). In contrast, only a minority of otoferlin particles (∼2.5%) from the nanodisc-bound dataset display alternative conformations of C_2_G (Fig. S2C). Altogether, these analyses suggest that the ‘closed’ state of otoferlin, where C_2_B, C_2_F and C_2_G are engaged in stable tertiary interfaces, is selected from an ensemble of sampled conformations in the lipid-free state, rather than being induced through membrane binding.

### Molecular dynamics simulations of the interaction of otoferlin with the target membrane

We cannot exclude that the other C_2_ domains interact with the target membrane *en route* to SV-docking or -fusion. In fact, the role of the C_2_C domain in preparing SVs for fusion (*26*) indicates that the domain might be required in interaction with the membrane and/or AZ proteins for a membrane-proximal SV to become fusion competent. Indeed, there is biochemical evidence of lipid binding by several C_2_ domains (e.g. ref. 25, 50). Moreover, it is possible that we missed lipid interactions of other C_2_ domains, e.g. in case they caused extensive remodeling and hence destabilization of the nanodisc membrane.

To gain more detailed insight into the nature and driving forces of the interactions between otoferlin and the membrane, we used atomistic molecular dynamics (MD) simulations. We used the membrane- and Ca^2+^-bound cryo-EM structure as a starting point and placed it so that its center of mass was ∼4.5 nm above the membrane. The membrane was modelled with CharmmGUI (*58*) and contained anionic lipids PS (30%) and PI(4,5)P_2_ (10%) mimicking the composition of the nanodisc. Otoferlin formed extensive contacts with the lipid membrane during the first microsecond of simulation (Fig. S5B). Internal distances between otoferlin residues differed by less than 5 Å from the original cryo-EM conformation, excluding disordered loops and the mobile C_2_G domain (Fig. S5A). Notably, the simulations reproduced the interactions between anionic lipids and the C_2_B, C_2_F, and C_2_G domains observed in the cryo-EM structure (Fig. 2). However, unlike in the cryo-EM structure, otoferlin formed a more extensive lipid interface in MD simulations involving all modelled C_2_ domains. Although all C_2_ domains were in contact with the membrane, most contacts were made by C_2_C and C-terminal domains (Fig. 5B). In contrast to the top loops of other C_2_ domains, C_2_C formed an amphiphilic helix which got submerged into the membrane (Fig. 5A). C_2_E formed a prominent membrane interface and strongly engaged with PI(4,5)P_2_ (Fig. 5E) which was mediated by the multiple basic residues in the top loops rather than by Ca^2+^ (Fig. 5D). C_2_D also engaged the membrane with its top loops, although the number of lipid contacts formed was lower than for the other Ca^2+^-binding C_2_ domains (Fig. 5B). Although C_2_D bound two Ca^2+^ ions, it did not bind PI(4,5)P_2_ or PS more often than phosphatidyl choline (PC, Fig. 5E). Notably, the Ca^2+^-binding top loops of C_2_D were pulled very close towards the membrane, rendering binding of large phosphorylated inositol headgroups of PI(4,5)P_2_ sterically unfavorable, while PI(4,5)P_2_ tended to cluster around the Ca^2+^-binding pockets of C_2_C, C_2_F, C_2_G domains (Fig. S5C). Moreover, different from C_2_E, the C_2_D top-loop periphery contains fewer basic residues which could form an interface with anionic lipids (Fig. 5C, D). Instead, a key contributor to the interaction with the lipid membrane in C_2_D appears to be the hydrophobic top loop, which inserts into the membrane (Fig. 5C).

**Figure 5.**
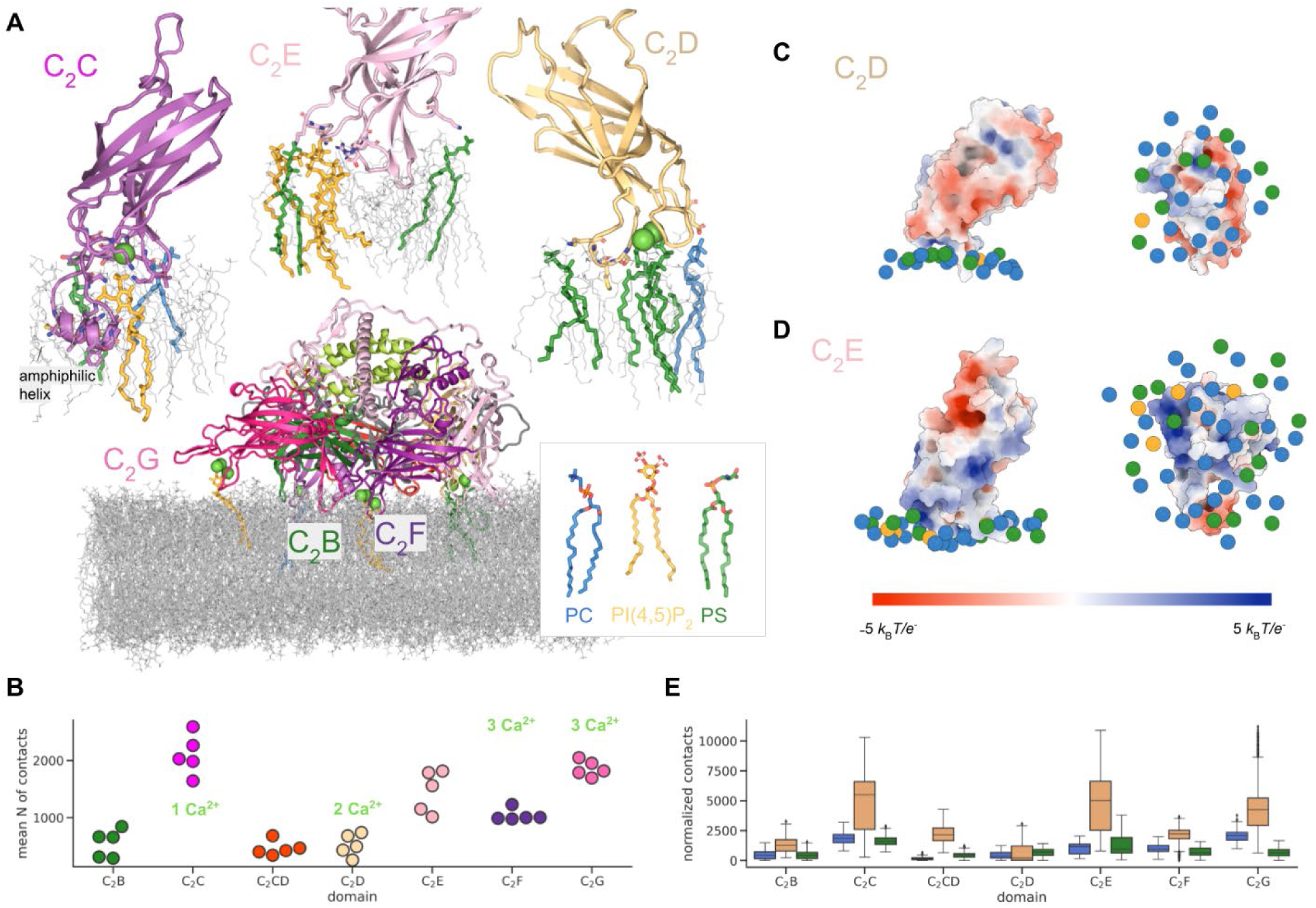
Molecular dynamics simulations reveal protein-lipid interaction driving membrane docking. **(A)** Overview of the multi-domain lipid interface observed in otoferlin simulations. Lipids within 3 Å of Ca^2+^ ions are highlighted in the overview image. C_2_C, C_2_D and C_2_E domains are shown enlarged in inserts. **(B)** Numbers of protein-lipid contacts established by individual C_2_ domains over the last 200 ns of simulations when the number of protein-lipid contacts plateaued (Fig. S5B). Mean values are shown for five simulation replicates indicated by individual dots. The number of calcium ions bound to domains is indicated for reference. **(C**, **D)** Electrostatic potential mapped onto accessible surface areas of C_2_D and C_2_E domains respectively. The potential was calculated with the APBS plugin in PyMol. Lipids within 15 nm of each domain are represented by their central phosphorus atom and colored according to the legend in A (also see Fig. S5C for lipid densities at the C_2_ domains). Side views are shown on the left, and bottom views are shown on the right. **(E)** Number of contacts between individual C_2_ domains and every lipid type in the last 200 ns of five simulation replicates normalized by the lipid abundance in the model membrane, colored according to the legend in A. The box boundaries represent the first and third quartiles of data distribution pooled together from five replicates, and the middle bar indicates the median value; the whiskers cover observations that are within 1.5 interquartile ranges of lower and upper quartiles; the rest of the data points are displayed individually.

To test how important Ca^2+^ ions and PI(4,5)P_2_ are for observed binding, we carried out two simulations of the membrane bound otoferlin where we removed either Ca^2+^ or the PI(4,5)P_2_ charges. After Ca^2+^-removal, the contacts between protein and membrane underwent a quick and long-term decrease, primarily because of detachment of C_2_G. In contrast, removing charges from the PI(4,5)P_2_ lipids did not have an immediate effect on membrane binding, and the number of lipid contacts was comparable to the unperturbed simulations (Fig. S5B, D). These findings are supported by biochemical data suggesting that Ca^2+^ plays a more important role than anionic lipids in liposome binding by otoferlin (Fig. S1G). Because neither of the perturbations caused rapid detachment of otoferlin from the membrane, we conclude that, once established, the otoferlin-lipid interface is reinforced and stabilized by hydrophobic interactions.

### Otoferlin facilitates SNARE-dependent membrane fusion and might mediate fusion itself

Our structural analyses of otoferlin revealed a complex membrane interaction mechanism, in which multiple C_2_ domain elements deeply insert into the lipid bilayer. Therefore, analogous to synaptotagmins, otoferlin may remodel and promote fusion between two lipid membranes in a Ca^2+^-dependent manner (*59–61*). To investigate otoferlin’s ability to promote vesicle docking and fusion *in vitro*, we established an ensemble lipid mixing assay (Fig. 6A, (*62*)). Guided by the mouse otoferlin (216–1931) cryo-EM structures, we expressed and purified a membrane-anchored otoferlin variant (residues 216-1992), which includes the transmembrane helix, but lacks the N-terminal C_2_A domain. Otoferlin (216–1992) was stable upon reconstitution into liposomes containing 5 mol% PS and retained its Ca^2+^-binding activity. As the SV fusion machinery at IHC synapses and the precise SNARE composition remain to be characterized (*9*, *11*, *13*, *14*) we chose to use well-characterized neuronal R- and Q-SNAREs for our assay, especially given their possible interactions with otoferlin (*6*). Notably, the precise SNAREs set does not appear to be critical for observing Ca^2+^- and C_2_ domain sensitive vesicle fusion, particularly when using minimized SNAREs constructs (*63*). Consequently, we co-reconstituted otoferlin (216–1992) with the rat R-SNARE VAMP2 (vesicle-associated membrane protein 2, also known as synaptobrevin-2) into large unilamellar vesicles (LUVs). These vesicles were then fused with a second LUV population containing the Qa-SNARE syntaxin-1A (rat, residues 183-288) and the Qbc-SNARE SNAP-25A (rat, residues 1-206) in the form of the ΔN-acceptor complex; in this complex, the Q-SNAREs are stabilized by a C-terminal VAMP2 fragment (residues 49-96, (*64*)). The acceptor ΔN liposomes were dual-labelled with the NBD-rhodamine FRET pair (*59*, *65*) and included acidic phospholipids (25 mol% PS or 25 mol% PS and 5 mol% PI(4,5)P_2_), closely resembling the nanodisc membranes used for cryo-EM structure determination.

**Figure 6.**
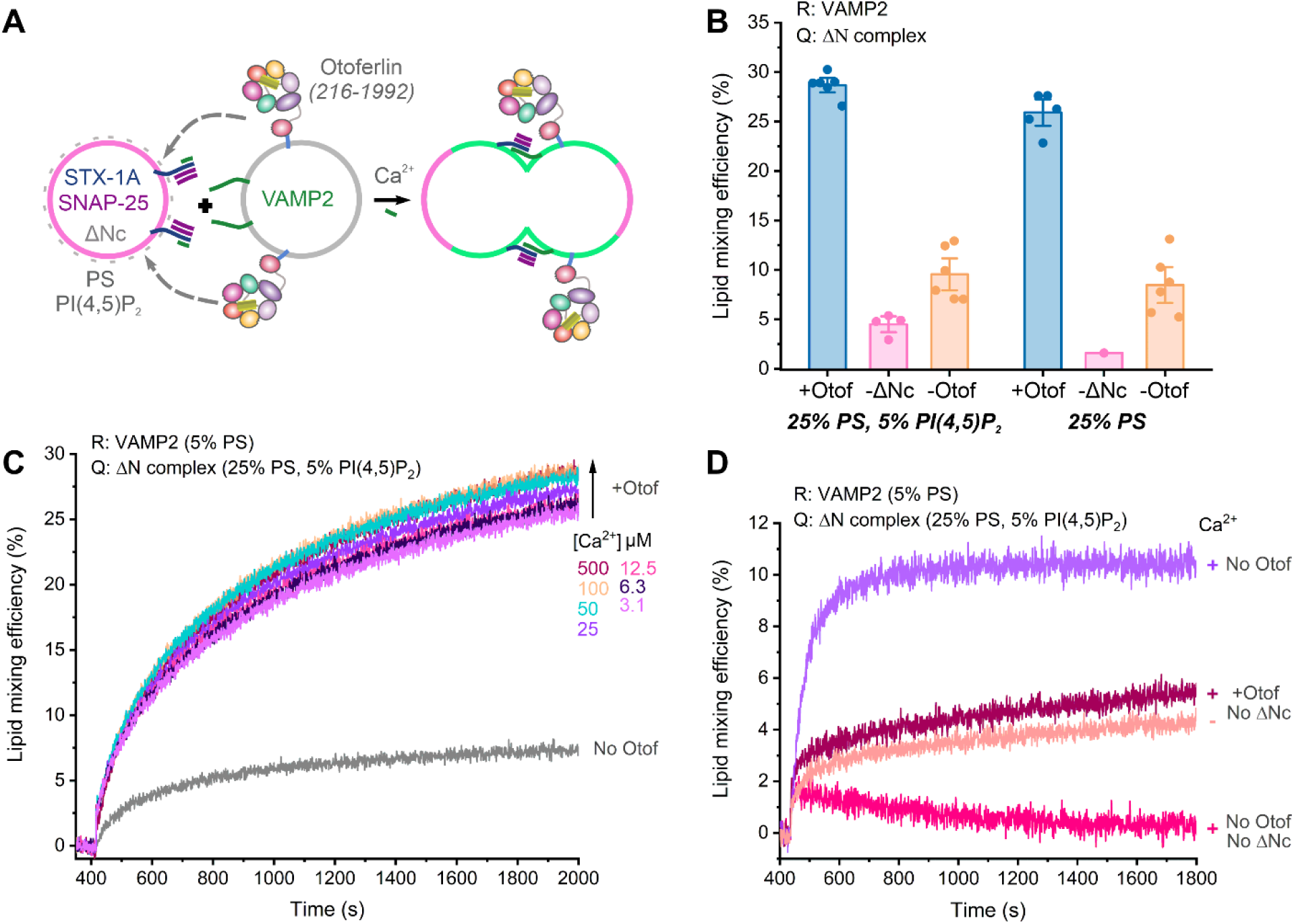
Otoferlin facilitates SNAREs mediated vesicle docking and fusion. (**A**) Schematic depicting the design of the otoferlin-sensitive ensemble lipid mixing assay. (**B**) Bar plot summarizing the final efficiencies of the otoferlin-sensitive liposome mixing reactions. The lipid mixing experiments were set up in the presence of 500 μM CaCl_2_ and the acceptor liposomes contained 25 mol% PS or 25 mol% PS and 5 mol% PI(4,5)P_2_. The final lipid mixing efficiencies were calculated after a 20 min incubation. The lipid mixing reactions were carried out either in the presence of membrane-anchored otoferlin, VAMP2, and Q-SNAREs (+Otof), with SNAREs-alone (-Otof), or without Q-SNAREs (ΔN complex) on acceptor liposomes (-ΔNc). Errors bars indicate the s.e.m. (**C**) Exemplary fluorescence traces of SNARE-dependent liposome mixing reactions in the presence (+Otof) or absence (No Otof) of membrane-anchored otoferlin (216-1992) at different [Ca^2+^]. In these reactions, otoferlin (216-1992) was co-reconstituted with VAMP2. Fluorescence traces of one representative experiment are shown. (**D**) Membrane-anchored otoferlin (216-1992) promotes SNARE-independent lipid mixing *in vitro*. Normalized fluorescence traces are shown for the reactions lacking otoferlin (No Otof), lacking Q-SNAREs (+Otof, No ΔNc), or lacking both (No Otof, No ΔNc). The lipid mixing experiments were carried out with 500 μM CaCl_2_ (+) or 500 µM MgCl_2_ (-).

Consistent with our cryo-EM structures, MD simulations, and supporting biochemistry, otoferlin (216–1992) accelerated SNARE-dependent liposome mixing in our assay, indicative of it facilitating vesicle docking for efficient membrane fusion. Importantly, we also observed increased lipid mixing efficiency in the presence of otoferlin compared to SNAREs alone (28.68±0.49% vs. 9.55±1.08%, Fig. 6B-C), when acceptor vesicles included PS or both PS and PI(4,5)P_2_. However, neither PI(4,5)P_2_ nor Ca^2+^ was essential for lipid mixing at physiological ionic strength (150 mM KCl, (*63*)), suggesting a strong Ca^2+^-independent component in otoferlin-dependent vesicle fusion, likely mediated by both hydrophobic and electrostatic interactions (Fig. 5). In contrast, stable liposome binding *in vitro* by soluble otoferlin (216–1931) was Ca^2+^-sensitive under similar buffer conditions (Fig. 3G and Fig. S1E). These assay differences suggest that partial membrane binding by otoferlin through Ca^2+^-independent mechanisms, without involving all C_2_ domains, may be sufficient to accelerate SNARE-dependent vesicle fusion *in vitro*. Such Ca^2+^-independent vesicle tethering or tight docking may involve the L3 loop of C_2_B, the L1 amphipathic helix of C_2_C, the basic surface of C_2_E, or hydrophobic residues in the top loops of C_2_F and C_2_G (Fig. 2 and Fig. 5). As these structural elements project from the same side of otoferlin, they may collectively engage the target vesicle, shorten the intermembrane distance, and thereby promote the nucleation of *trans*-SNARE complexes – potentially without directly interacting with SNAREs. Intriguingly, low efficiency lipid mixing was observed in the absence of the ΔN-acceptor complex (efficiency of 4.5±0.54%), indicating that otoferlin alone can promote vesicle docking and fusion *in vitro* (Fig. 6D). In such a mechanism, through deep insertion at several sites, otoferlin would promote local membrane remodeling and curvature changes, which, combined, may lower the energy barriers for lipid bilayer fusion (*63*) (Fig. 2, Fig. 4C, and Fig. 5A). Future studies will be needed to test the physiological relevance of this mode of otoferlin action for SV fusion events at IHC synapses.

### *In vivo* analysis of otoferlin function

We next turned to *in vivo* analysis of otoferlin function to test the ‘Ca^2+^-sensor of SV-fusion’ hypothesis. Following efforts targeting C_2_C (*26*), C_2_F (*29*) and C_2_G (*36*) domains that had supported but not unequivocally demonstrated such a role, we performed CRISPR/Cas9 mutagenesis of the C_2_D domain to alter Ca^2+^-binding by otoferlin in mice. Our structure-guided mutagenesis aimed to progressively disrupt Ca^2+^-binding to the C_2_D domain: substitution of aspartates D1060, D990, and D990/996 by alanine is predicted to alter the binding of one (*Otof^D1060A^* and *Otof^D990A^*) or both (*Otof^D990-996A^*) Ca^2+^ ions (Fig. 3). As *Otof* mutations often lead to decreased otoferlin protein levels, alter the subcellular otoferlin distribution, or cause a reduction in the number of afferent IHC-SGN synapses (*31*, *32*, *36*), we first performed immunohistochemistry in the organ of Corti early after hearing onset (2-weeks-old) and after full cochlear maturation (5-8-weeks-old). Immunofluorescence of the SV-marker Vglut3 (*7*, *66*) was unaltered in IHC of the mouse mutants and served as a reference for the subcellular distribution of otoferlin (Fig. 7A-E, Fig. S6, S7). We found the levels and subcellular distribution of otoferlin to be unaltered in IHCs of homozygous *Otof^D990-996A^* and *Otof^D1060A^* mice, while homozygous *Otof^D990A^* IHCs exhibited a mild reduction to 71.10±0.03% of WT levels, but unaltered subcellular otoferlin distribution (Fig. 7D, E). The number of IHC-SGN synapses, quantified by counting juxtaposed spots of presynaptic RIBEYE/CtBP2 and postsynaptic Homer1 immunofluorescence, was normal in all three homozygous mutants (Fig. S7).

**Figure 7.**
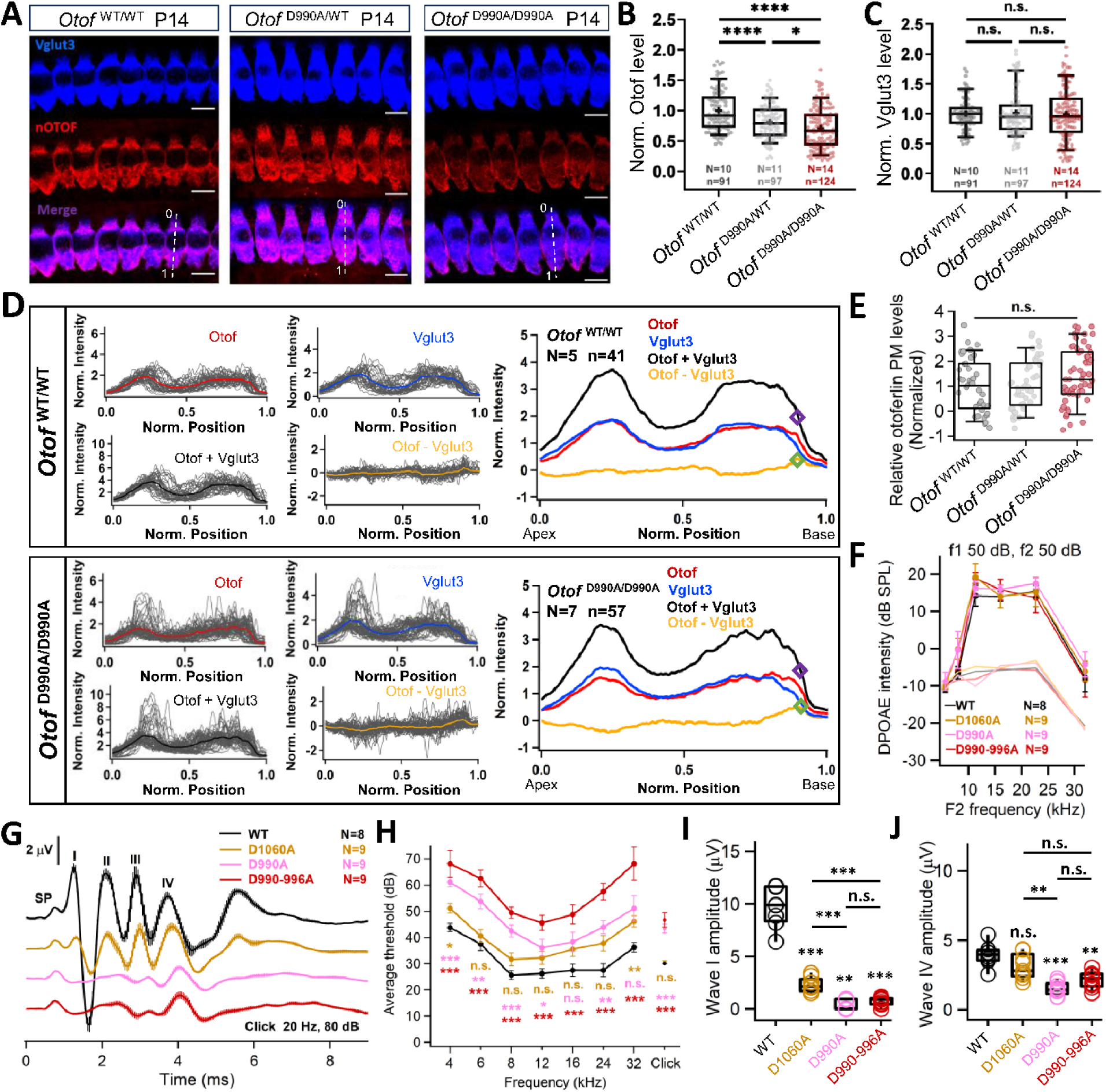
Disruption of Ca^2+^-binding by the C_2_D domain of otoferlin impairs hearing. **(A)** Staining IHCs with otoferlin (OTOF) and Vglut3 antibodies. White dashed lines indicate the line scans in (D), with 0 and 1 indicating start- and endpoints. Scale bar, 10 μm. **(B)** Otoferlin immunofluorescence intensity indicated reduced level (0.711±0.031) in *Otof^D990A^* IHCs compared to littermate control IHCs **(C)** Vglut3 immunofluorescence intensity indicated unchanged Vglut3 levels in *Otof^D990A^* IHCs compared to littermate control IHCs. **(D)** Line profile analysis. For quantification of membrane staining, the fluorescence was normalized to the cellular fluorescence for each fluorophore. The sum of both fluorescence values (black line) was used to determine the position of the basal membrane. At the most basal cellular point along this line which exceeds the threshold value of 2 (purple diamond), the otoferlin-Vglut3 fluorescence difference (yellow line) gave the value for relative otoferlin plasma membrane levels (green diamond). **(E)** Relative levels of membrane-bound otoferlin at the basal pole of IHCs. **(F)** Robust DPOAE in homozygous *Otof^D1060A^* (yellow), *Otof^D990A^* (magenta) and *Otof^D990-996A^* (red) mice indicate normal OHC function across the cochlear frequency range. DP-gram shows amplitude for different pairs of primary tones for different frequency pairs. **(G)** Auditory brainstem responses (ABR) of *Otof^wt^* (black), homozygous *Otof^D1060A^* (yellow), *Otof^D990A^* (magenta) and *Otof^D990-996A^* (red) mice to 80 dB click stimulation. **(H)** ABR thresholds, **(I)** ABR wave I amplitude and **(J)** ABR wave IV amplitude (same color code as in A). Box and whisker plots represent median, 25th and 75th as well as 10th and 90th percentiles.

*Otof^D1060A^, Otof^D990A^*, and *Otof^D990-996A^* mice showed significantly impaired synaptic transmission at IHC-SGN ribbon synapses of increasing severity as demonstrated by *in vivo* and *ex vivo* physiology. Importantly, cochlear amplification mediated by outer hair cells was intact as evident by otoacoustic emissions (Fig. 7F, S8), supporting the notion of an auditory synaptopathy (*15*). Recordings of auditory brainstem responses (ABRs, Fig. 7G) showed a mild (∼20dB sound pressure level, SPL) but significant increase of sound threshold for 2 months-old homozygous *Otof^D990A^* and *Otof^D990-996A^*mice (Fig. 7H) as compared to pooled littermate wild-type controls (*Otof^wt^*). Homozygous 2 months-old *Otof^D1060A^* mice showed a non-significant trend toward higher ABR thresholds (Fig. 7H). Synchronous activation of the SGN population, approximated as amplitude of ABR wave I, was significantly impaired in all mutants. Wave I amplitudes declined with the anticipated strength of disrupting C_2_D Ca^2+^-binding (Fig. 7I). Interestingly, central ABR waves were less affected (Fig. 7G, J), which likely reflects improved neural synchrony due to convergent input of SGNs into the cochlear nucleus as previously reported for mouse mutants with impaired synaptic sound encoding (*24*, *32*, *34*, *67*).

Next, we turned to recordings from individual SGNs to scrutinize the impact of impaired Ca^2+^-binding on spontaneous and sound evoked synaptic transmission from IHCs *in vivo.* Spontaneous SGN firing, reporting transmission at the IHC resting potential, tended to be lower in *Otof^D990A^* and *Otof^D990-996A^* but not in *Otof^D1060A^* mutants (Fig. S9A). We note that we might have underestimated the effect of the *Otof* mutations on spontaneous firing by the use of isoflurane anaesthesia. Isoflurane causes lower spontaneous firing rates (*68*) likely due to inhibition of Ca_V_1.3 channels of IHCs (*69*). As expected, given intact cochlear amplification, we found thresholds and frequency tuning to be unaltered in SGNs of all three mutants (Fig. S9B, C). We then studied the SGN responses to suprathreshold tonebursts (30 dB above threshold, repetition rate of 5 Hz), which allows one to study release probability and readily releasable vesicle pool (RRP) dynamics of IHC AZs (*70*) (Fig. 8A). Most SGNs in the mouse cochlea receive input from a single AZ (*71*) and their SGN firing rate reports the rate of SV release convolved with neural refractoriness (*72–74*). Assuming unaltered SGN excitability in the *Otof* mutants, we can use onset firing rate, first spike latency and its variance as well as short-term spike rate adaptation to comparatively assess the rate of initial SV release as well as kinetics of RRP depletion, respectively (*73*). Initial SV release is co-determined by release probability and the size of RRP, while the RRP depletion time constant is governed by the rate constants of release (i.e. reporting SV release probability) and of SV replenishment (*70*). The adapted firing rate reflects sustained release which is governed by balanced SV fusion and replenishment and also requires sustained Ca^2+^-influx (*70*). *Otof* missense mutants studied at the single SGN level to date showed strong firing rate adaptation lending support to a critical role of otoferlin in SV replenishment (*31*, *32*). Interestingly, initial firing was near normal at low stimulation rate (2Hz) in the mouse model of the human p.Ile515Thr missense mutation (*32*).

**Figure 8.**
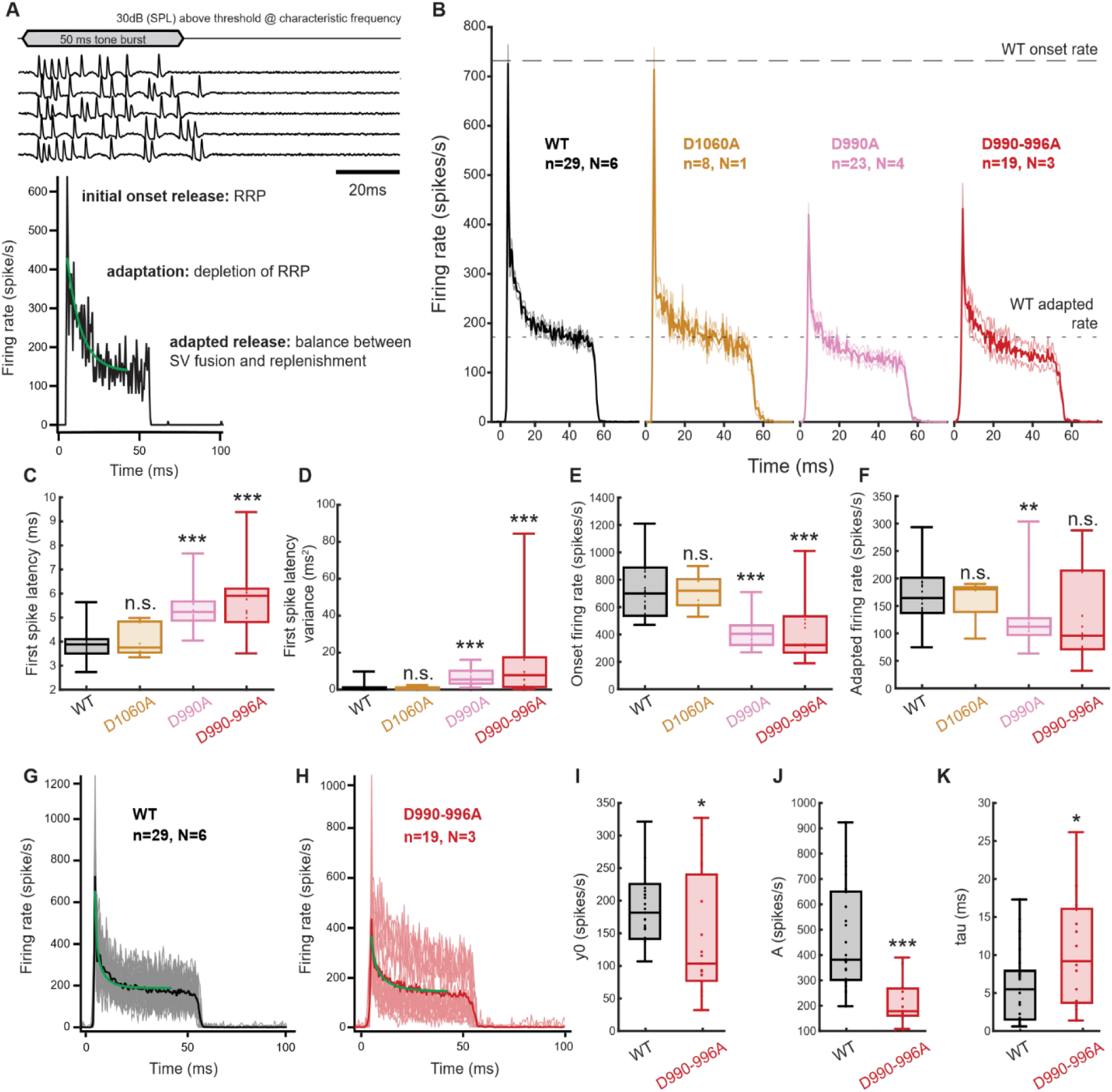
Impaired IHC-SGN synaptic transmission diminishes SGN onset firing and spike rate adaptation, indicating reduced IHC synaptic vesicle release probability. **(A)** Example SGN spike traces (top panel) recorded in response to 50 ms tone bursts at characteristic frequency, 30 dB (SPL) above threshold and resulting peristimulus time histogram (PSTH) of the SGN firing rates across 200 repetitions (bottom panel). Single exponential fitting of the spike rate adaptation is shown in green. Onset firing rate is dependent on initial release from the standing readily releasable pool (RRP) while adaptation to the adapted firing rate depends on the balance between RRP depletion and vesicle replenishment. **(B)** PSTHs of SGN firing rates recorded as depicted in (A) for *Otof^w^*^t^ (black, N = 6, n = 29), homozygous *Otof^D1060A^* (yellow N =1, n = 8), *Otof^D990A^* (magenta, N = 4, n = 23), and *Otof^D990A-996A^* (red, N = 3, n = 19) mice. Mean traces are displayed with SEM indicated by fainter lines. **(C)** Median first spike latency, **(D)** first spike latency variance, **(E)** onset firing rate, and **(F)** adapted firing rate of the SGNs shown in (B). **(G-K)** Single exponential fitting of individual PSTHs of **(G)** Otof^wt^ and **(H)** *Otof^D990-996A^* SGNs for which average traces are displayed (average traces are also shown B) with the averaged fits shown in green. Individual PSTH traces are shown as fainter lines. We note that the spike rate adaptation kinetics reflects both neural refractoriness and short-term synaptic depression (*e.g.* 24) which can be more precisely described by fitting a double exponential function or a model to it. On the assumption that the impact of refractoriness will be comparable for SGNs of both genotypes we chose the simplest fit function to approximate the kinetics of RRP depletion (K) for comparing release probability. **(I)** The plateau of the fit, y0, **(K)** the amplitude of the adapting component, and **(J)** the time constant of adaptation, tau for the fitted PSTHs shown in (G) and (H). Fitting was done in Igor Pro. Data in (C-F, I-K) are shown as box and whisker plots with data points overlaid, median, 25th and 75^th^ percentiles (box), 10th and 90th percentiles (whiskers) displayed.

Onset firing was significantly reduced and first spike latency as well as its variance were increased in *Otof^D990A^*and *Otof^D990-996A^* mutants, but not in *Otof^D1060A^*mutants, which indicates a reduced initial SV release rate at *Otof^D990A^*and *Otof^D990-996A^* AZs (Fig. 8B-E). Exponential fitting to the sound-evoked firing rate revealed slower kinetics of spike rate adaptation in *Otof^D990-996A^*mutants (Fig. 8G, H, K) which indicates a reduction of SV release probability and/or of SV replenishment. Steady state release, reported by the adapted firing rate, was not significantly changed in *Otof^D990-996A^* and *Otof^D1060A^*mutants, contrasting the *Otof^I515T^* mutant and indicating largely intact SV replenishment during ongoing stimulation (Fig. 8F, but smaller in *Otof^D990A^*, p= 0.001). Therefore, we propose that reduced release probability, rather than a reduced RRP size due to impaired SV replenishment, is the primary cause of the sound encoding deficit that includes a diminished amplitude of the adapting component (Fig. 8J) and trend to lower adapted firing rate of *Otof^D990_996A^*SGNs. This notion was further supported by the fact that onset and steady state firing rate of all three mutants did not change with the stimulation rate beyond what we found in wild-type SGNs (10 and 0.5 Hz, Fig. S9). This again sets apart the C_2_D mutants from other *Otof* mutants (*31*, *32*). In summary, our *in vivo* analysis suggests that the altered Ca^2+^-binding of otoferlin impairs sound encoding by reducing release probability.

To directly assess Ca^2+^-triggered IHC-exocytosis, we turned to perforated-patch recordings of exocytic membrane capacitance changes (ΔC_m_) in response to depolarization that elicits maximal Ca^2+^-influx (to −17 mV) from IHCs of all three *Otof* mutants (Fig. 9A-O). Ca^2+^-influx was not significantly altered in voltage-dependence and amplitude (Fig. 9, Fig. S10). We found a significant reduction of ΔC_m_ elicited by brief (20 ms) depolarizations that tap initial release from the RRP in *Otof^D990A^*(3.35±0.96 fF vs. 7.77±1.01 fF for littermate control, p=0.0047, Fig. 9H) and *Otof^D990-996A^* (3.29±0.70 fF vs. 8.36±0.74 fF for littermate control, p = 0.00007, Fig. 9M) IHCs as compared to wild-type IHCs obtained from littermates.

**Figure 9.**
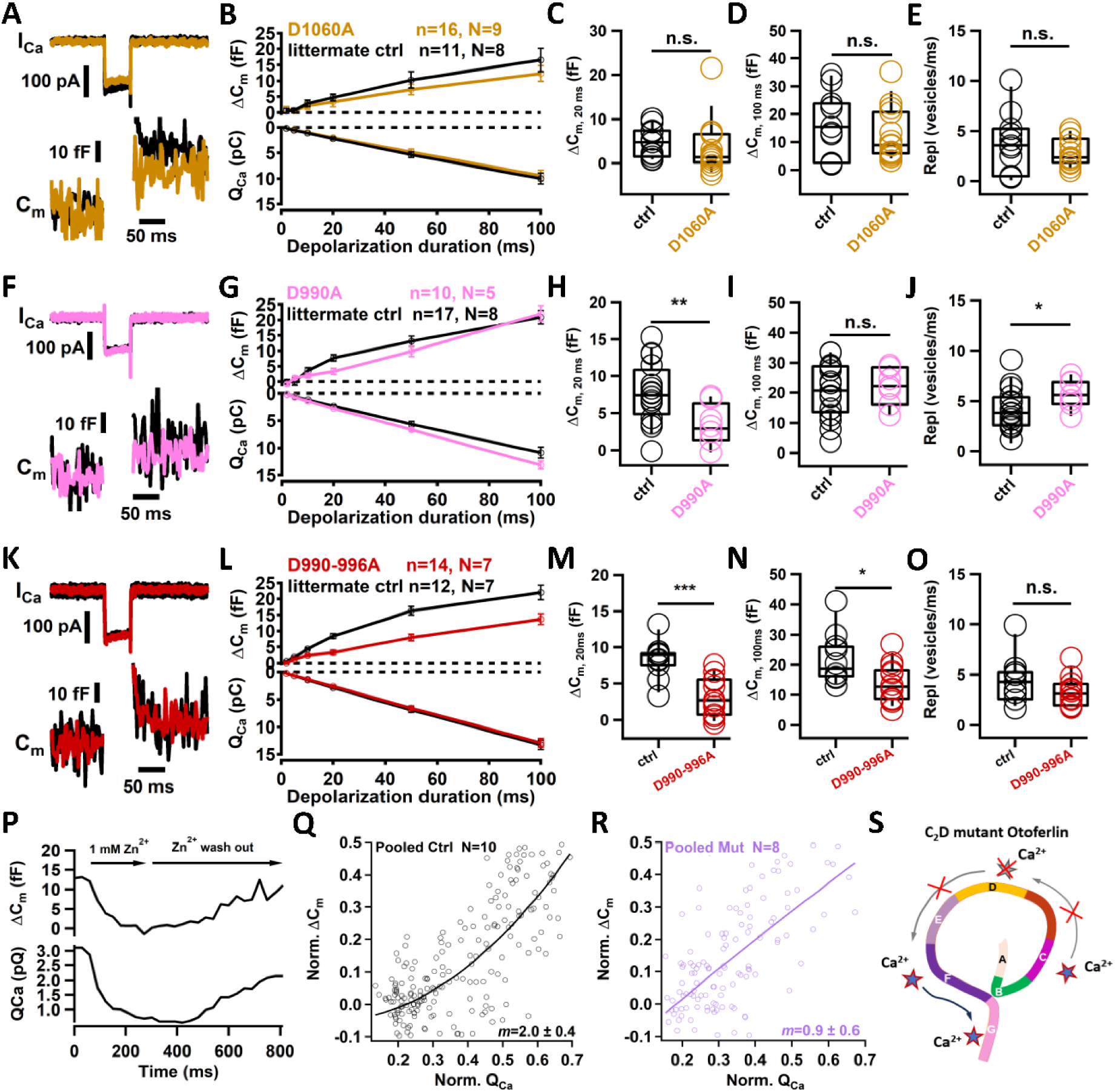
Impaired synchronous exocytosis and linearization of the Ca^2+^-dependence of IHC exocytosis. **(A, F, K)** Exemplary membrane capacitance changes (ΔC_m_, upper) and Ca^2+^ currents (lower) elicited by 50 ms depolarizations to −17 mV in IHCs of *Otof^wt^* (black), homozygous *Otof^D1060A^* (yellow), *Otof^D990A^* (magenta), and *Otof^D990-996A^* (red) mice. **(B, G, L)** ΔC_m_ (upper) and integrated Ca^2+^ current integral (Q_Ca_,lower) elicited by depolarizations to - 17 mV for different durations in IHCs, same color code and numbers of IHCs and mice as in **(A, F, K)**. **(C, H, M)** RRP exocytosis reported by ΔC_m,20ms_ (20 ms depolarizations to −17 mV), same color code and numbers of IHCs and mice as in **(A, F, K)**. **(D, I, N)** Exocytosis reported by ΔC_m,100ms_ (100 ms depolarizations to −17 mV), same color code and numbers of IHCs and mice as in **(A, F, K)**. **(E, J, O)** Maximal rate of sustained exocytosis (ΔC_m_ per ms during 20 and 100 ms of depolarization to −17 mV), same color code and numbers of IHCs and mice as in **(A, F, K)**. **(P)** Exemplary estimation of the apparent Ca^2+^-dependence using rapid flicker blocking by 1 mM Zn^2+^ in a control IHC: ΔC_m_ (upper) and Q_Ca_ (lower). **(Q and R)** Scatter plots ΔC_m_ vs. Q_Ca_ of control (q) and mutant (pooled *Otof^D1060A^* (n=3) and *Otof^D990A^* (n=5)) and power function fits to the data. **(S)** Suggested sequential Ca^2+^-binding initiated by C_2_C binding and proceeding to the C-terminus, which is disrupted in the C_2_D mutants leading to a linear Ca^2+^-dependence. Data in **(A, D)** are shown as mean SEM, box and whisker plots with data points overlaid, show median, mean, 25^th^ and 75^th^ percentiles (box), 10^th^ and 90^th^ percentiles (whiskers). Two sample comparisons were performed using t-test or the Wilcoxon rank test. Significant differences are indicated as *P < 0.05, **P < 0.01, ***P < 0.001.

ΔC_m_ in response to longer stimuli in *Otof^D990A^* IHCs were smaller than those of littermate controls, while those of *Otof^D990-996A^*and *Otof^D1060A^* IHCs did not differ from their controls (Fig. 9D, I, N). Importantly, we did not find indication for reduced SV replenishment when analysing the rate of sustained exocytosis (SVs/ms between 20 and 100 ms of stimulation, assuming a SV capacitance of 40 aF, (*75*), (*Otof^D990A^*: 5.75±0.55 SVs/ms vs. littermate control: 4.04±0.56 SVs/s, p=0.04235 and *Otof^D990-996A^*: 3.24±0.37 SV/ms vs. littermate control: 4.42±0.68 SVs/ms, p=0.14544, Fig. 9E, J, O). This is consistent with the largely unaltered adapted SGN firing rates *in vivo*.

Finally, we examined the apparent Ca^2+^-dependence of RRP exocytosis of *Otof^D990A^* and *Otof^D1060A^* IHCs when gradually reducing the Ca^2+^-influx elicited by 20 ms depolarizations (to −17 mV) by slow bath perfusion of Zn^2+^ (Fig. 9P) (*20*, *22*), which causes a rapidly flickering channel block (*76*). Fitting the relationship of normalized ΔC_m_ (≤50%, to avoid saturation of the function by RRP depletion) and the integrated normalized Ca^2+^-influx (Q_Ca_, 70%) by a power function we observed that the Ca^2+^-cooperativity of RRP exocytosis dropped to a power *m* of 0.9±0.6 compared to 2.0±0.4 for *Otof^wt^* IHCs (Fig. 9Q, R). We postulate that the disrupted Ca^2+^-binding of C_2_D prohibits the progress of Ca^2+^- and phospholipid-dependent membrane interaction of otoferlin (Figs. 3-5) *en route* to SV fusion (Fig. 10).

**Figure 10.**
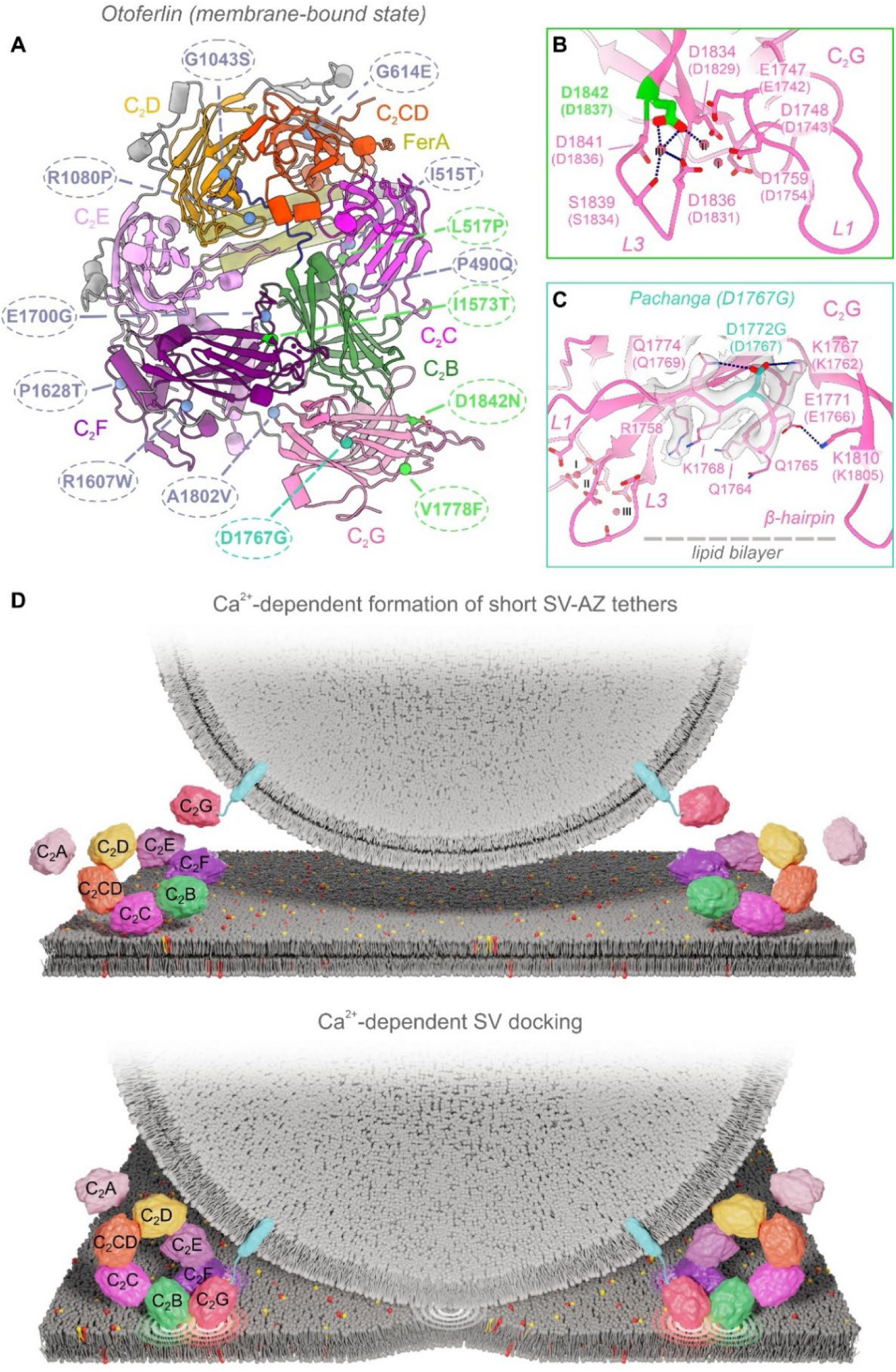
Predicted impact of *OTOF* mutations and model of otoferlin-mediated SV docking. **(A)** Selected *OTOF/Otof* missense mutations with amino acid substitutions mapped onto the membrane bound structure of human otoferlin. Amino acids in mouse sequence are given in parentheses. Dark blue font indicates mild hearing impairment, green font moderate or severe hearing impairment (Table S2). (**B** and **C**) Structural prediction of mutations affecting C_2_G. **(B)**: D1842 human missense mutation: D1842N and **(C)**: mouse mutants (D1837A) functionally likely equivalent to D1842N (B) and (D1767G). **(D)** Model of otoferlin-mediated SV docking inspired by structural and functional data.

## Discussion

We present a structure-function analysis of otoferlin, a deafness gene product, for which the first gene replacement therapy of the inner ear is currently in clinical trials. High resolution cryo-EM structures of lipid-free and membrane-bound soluble otoferlin and molecular dynamics simulations inform a model of membrane interactions which we propose to mediate tethering and docking of SVs (Fig. 10). The structures and the corresponding model provide a basis for interpretation of human *OTOF* mutations (Fig. 10A-C). Towards deciphering Ca^2+^-triggered SV fusion at IHC synapses, we elucidated the structural basis of Ca^2+^-binding by otoferlin and tested the ‘Ca^2+^-sensor of SV-fusion hypothesis’ by multiscale physiology in mouse mutants. Specifically, we show that C_2_C, C_2_D, C_2_F, and C_2_G bind Ca^2+^ by their top-loops but differ in the number of Ca^2+^ ions coordinated and the contribution of anionic phospholipids. In particular, the membrane bound otoferlin structure indicates that C_2_C-C_2_D can coordinate Ca^2+^ without the contribution of anionic phospholipids. Structure-informed progressive alteration of Ca^2+^-binding by the C_2_D domain of otoferlin led to increasing deficits of synaptic sound encoding. Mechanistically, we observed a reduced release probability and a loss of Ca^2+^-cooperativity of RRP exocytosis. This is reminiscent of the findings in hippocampal neurons of mouse mutants with impaired Ca^2+^-binding by synaptotagmin 1 (*27*) and verifies the hypothesis that otoferlin is the Ca^2+^-sensor of SV-fusion in IHCs. Different from a previous analysis of *Otof^D515-517A^* mice (*26*) with altered Ca^2+^-binding of C_2_C, we found SV-replenishment to be largely unaltered. This allowed us to isolate the ‘Ca^2+^-sensor of SV-fusion function’ of otoferlin and to conclude that Ca^2+^-binding to C_2_D is dispensable for Ca^2+^-dependent SV-replenishment. We propose that Ca^2+^-binding to the C_2_D domain triggers the interaction with C_2_E domain involving top loop 1 of C_2_D and top loop 4 of C_2_E. The resulting C_2_E domain rotation might facilitate the Ca^2+^- and phospholipid-dependent interaction of the C_2_F, C_2_G with C_2_B and the target membrane. It is possible that this sequence of Ca^2+^-binding steps as a whole is reflected in the supralinear Ca^2+^-dependence of SV exocytosis in IHCs (*20*, *22*, *30*). Our data is also consistent with the possibility that Ca^2+^-bound otoferlin directly mediates SV-fusion.

A role of otoferlin in preparing SVs for fusion has been proposed based on ultrastructural and physiological mouse analyses in mutant mice, and our structural data now provide a basis for interpretation. Specifically, electron tomography of IHC synapses of *Otof^-/-^* mice revealed a lack of short tethers (<20 nm) connecting membrane-proximal SVs to the AZ membrane. These short ‘tethers’ are well compatible with the dimensions of SV-standing otoferlins (∼10 nm) that might mediate initial contacts with the AZ membrane via its top-arch, e.g. by C_2_C domain (model in Fig. 10) which showed extensive lipid interactions in MD simulations (Fig. 5).

We postulate that SVs then proceed to docking by progressive membrane interaction of otoferlin. The *Otof^D1772G^* mutation (D1767G in mice) carried by *Pachanga* mouse mutants (*31*, *77*) likely modulates the network of hydrogen bonds between the C_2_G core and the C_2_G beta-hairpin (depicted in Fig. 10A, C). We propose that the mutation affects the tight C_2_G interaction with the membrane and the interface of C_2_G with C_2_F. While the Ca^2+^-mediated electrostatic interactions of the C_2_G top loops with the membrane might allow for SV docking, a tighter hydrophobic contact between C_2_G and the membrane could be required for tight docking to gain full fusion competence of the SV. Consistent with such a scenario, electron tomography of stimulated IHC AZs of *Pachanga* mice showed an increase of multiply tethered and docked SVs (*78*).

The otoferlin structure predicts the human *OTOF* variant p.D1842N (*53*) (Fig. 10A, B and Table S2) to disrupt the binding of two Ca^2+^ by the C_2_G top loops, which we interpret to interfere with SV docking. Mouse mutagenesis targeting C_2_G Ca^2+^-binding (*Otof^DDA^,* pD1836/1837A in mouse, p.D1841/1842A in human) revealed deafness due to a complete loss of IHC exocytosis despite substantial (∼60%) basolateral otoferlin levels (*36*). Altered subcellular distribution with increased apical and decreased basolateral otoferlin and block of exocytosis were taken to indicate impaired otoferlin membrane targeting and a critical role of C_2_G Ca^2+^-binding.

The precise sequence of Ca^2+^- and lipid binding events and accompanying conformational changes in otoferlin remain to be investigated by further cryo-EM, MD simulations and potentially MINFLUX (*79*). We hypothesize that SVs get attracted to the AZ membrane via otoferlin-independent longer range tethering mechanisms (*12*) and proceed to short-range tethering and docking upon progressive engagement of otoferlin with the AZ membrane (Fig. 10D). Otoferlin might sample the open and closed formation and upon contacting the membrane via SV-distal C_2_C-C_2_E domains transition (Fig. 10D) to the closed configuration pulling the SV and AZ membranes closer to each other by the SV-proximal C_2_F and C_2_G engaging with the AZ membrane (Fig. 10D).

### Limitations of the study

Full elucidation of the molecular mechanisms underlying otoferlin-dependent SV-fusion will require integration of further molecular, structural and functional data. For example, deciphering the molecular events underlying membrane fusion will require future work. Questions remaining include: i) How reversible are the tethering and docking reactions? ii) Which proteins compose the fusion machinery, what is their stoichiometry and how many copies are required?

## Acknowledgments

We thank Christiane Senger-Freitag, Sandra Gerke, Sina Langer, Andy Schacht, and Patricia Räke-Kügler for excellent technical and administrative support. We are grateful to Drs. Tat Cheung Cheng and Ruben Fernandez-Busnadiego for supporting cryo-EM data acquisition and Dr. Eri Sakata for critically reading the manuscript.

## Author contributions

Conceptualization, T.M., H.C., C.C., and J.P. Methodology, N.Ba., H.G., F.L., T.M., C.P., J.P., N.S., and C.W., Investigation, H.C., C.C., A.T., N.E., N.Ba., L.F., F.L., F.B., S.M., K.E., and C.W.; writing—original draft, T.M., H.C., C.C., and N.E; writing—review & editing, all authors; funding acquisition, N.Br., H.G., T.M., C.P., J.P., N.S., and C.W.; resources, N.Br., H.G., T.M., C.P., J.P., N.S., and C.W.; supervision, N.Br., H.G., F.L., T.M., C.P., J.P., N.S., and C.W.

